# Dichotomy of cellular inhibition by small-molecule inhibitors revealed by single-cell analysis

**DOI:** 10.1101/038000

**Authors:** Robert Vogel, Amir Erez, Grégoire Altan-Bonnet

**Affiliations:** ImmunoDynamics Group, Program in Computational Biology and Immunology, Memorial Sloan Kettering Cancer Center, 1275 York Avenue, Box 460, New York, N.Y. 10065, USA.; ImmunoDynamics Group Current Address, Cancer … Inammation Program, Center for Cancer Research - National Cancer Institute, Bldg 37 - Room 4134B, 37 Convent Drive - Bethesda MD 20892, USA.

**Author notes:** E–mail (Grégoire Altan-Bonnet).

## Abstract

Despite progress in developing small molecule inhibitors, a quantitative understanding of drug action in the physiological context of cells is lacking. Here, we apply single-cell analysis of signal transduction and proliferation to probe cellular responses to small molecule inhibitors. We use the model of cellular activation of T lymphocytes responding to cytokines and antigens. We uncover two distinct modes of drug action, in terms of signaling inhibition: digital inhibition (*e.g*. when the fraction of activated cells diminishes upon drug treatment, but cells remaining active appear unperturbed), and analog inhibition (*e.g*. when the fraction of activated cells is unperturbed while their overall activation is diminished). We introduce a computational model of the signaling cascade in order to account for such dichotomy. We test the predictions of our model in terms of the phenotypic variability of cellular responses under drug inhibition. Finally, we demonstrate that the digital/analog dichotomy of cellular response as revealed on short timescales with signal transduction, translates into similar dichotomy on long timescales. Overall, our analysis of drug action at the single cell level illustrates the strength of quantitative approaches to translate the promise of *in vitro* pharmacology into functionally-relevant cellular settings.

## Introduction

Individual cells rely on biochemical signaling pathways to translate environmental cues into physiological responses. Spurious activation of these pathways results in a cell’s mischaracterization of environmental conditions and aberrant cellular behavior. This behavior can, in some cases, be detrimental to the health of the organism - causing ailments such as inflammatory diseases (*e.g*. ulcerative colitis [1]), auto-immune disorders [2, 3] and cancer [4]. Inhibiting specific dysfunctional components with small-molecule chemical inhibitors has been successful in reducing aberrant signals and ultimately ailments [5]. Examples include Imatinib in treating Chronic Myelogenous Leukemia [6] and Gefitnib for patients with EGFR mutant nonsmall-cell lung cancer [7, 8]. However, despite these successes, inhibitory drug development remains slow and can benefit from new techniques to aid screening of candidate compounds [9, 10].

Fundamentally, an effective chemical inhibitor acts on a signaling pathway by binding to the targeted enzyme and shutting down its enzymatic activity. In this context, optimizing a drug inhibitor abounds to optimizing its specific binding to the enzyme target of choice. Recent technological advances have focused efforts to development of pipelines that characterize drug specificity with respect to all human kinases *in vitro* [11, 12, 13] and in cell lines as models for physiological settings [14]. The emphasis on protein kinases is due to their prominence in signal transduction pathways, where they serve as information relays by transferring a phosphate from ATP to their target substrate. The technological advances in drug screening often fall short of anticipating the downstream consequences of drug inhibition: whereas kinase inhibition is optimized at the local (molecular) level, the response at the level of the entire pathway often remains sub-optimal. Consequently, it is difficult to predict cellular response to chemical inhibition. To gain some understanding of this response, emphasis has been placed on high throughput characterization of the response of cell lines [15, 16].

Despite these advances in drug screening, many poorly performing compounds proceed to, and often fail at, the organismal stage of drug discovery. This suggests that we may need to re-evaluate the relevance of bulk measurements on cell line models to drug development, emphasizing instead a more mechanistic understanding of individual primary cell responses to inhibition ( *“Whats wrong with drug screening today”* [17]). This need has been partially addressed by pioneering studies that characterized biochemical networks of primary cells [18, 19] and canonical cell type responses to inhibition [20, 21, 22]. Yet, while these studies have been illuminating, mechanistic principles of cellular responses to small-molecule chemical inhibition have remained elusive. It is precisely this gap in knowledge that this *Communication* attempts to address.

We conjecture that one needs to resolve diverse enzymatic states (*e.g*. phospho-status) at the single cell level in order to identify the complex nonlinear responses of signaling networks to inhibition. Nonlinear responses are often dominated by a subset of enzymes that determine the behavior of the pathway. Identifying these key enzymes uncovers novel vulnerabilities of the signaling network to inhibition [23]. Examples of nonlinear responses uncovered by single cell measurements are numerous: flow cytometry measurements of double-phosphorylated ERK (ppERK) accumulation in stimulated T lymphocytes exhibit a highly nonlinear bimodal response to antigen [24]; by imaging ERK in live cells, individual cell response to growth factors was shown to be pulsatile [25]; administration of either the tyrosine kinase inhibitor Gefitnib or the MEK inhibitor PD325901 yielded either a frequency or a mean reduction in ppERK signaling, respectively [26]. Similarly, epidermal-growth-factor (EGF) stimulation of 184A1 cells, a mammary epithelial cell line, revealed oscillatory ERK nuclear localization with a period invariant to EGF dose; at the same time the number of cells exhibiting oscillatory behavior reduced markedly with increased cell density and decreased EGF concentration [27]. These are but few examples of the dynamic complexities of biochemical signaling networks under stimulation, as revealed by single-cell measurements. In all these examples, by directly revealing different *modes* of inhibition, inaccessible by “high-throughput” population level measurements, single-cell measurements added crucial understanding to the structure of a signaling pathway.

In this study, we integrate single cell multi-parametric phospho-flow cytometry measurements, Cell to Cell Variability Analysis (CCVA, [28, 29]), and mechanistic modeling to dissect the mechanism of action of kinase inhibitors in primary mammalian cells. We provide both experimental and theoretical evidence that the network structure, in which the targeted enzyme is embedded, determines the signaling response to inhibition. Furthermore, we investigate the influence of protein expression variability on the sensitivity of single cells to inhibition. Following up on these insights we present experimental results demonstrating the functional relevance of our model of drug inhibition to cell proliferation, thereby bridging the short molecular timescale with the longer functional one. Taken together, in this *Communication* we demonstrate how, by combining mechanistic models and single cell measurements of primary cells, it is possible to predict cellular behavior in response to targeted molecular inhibition. We present a general and easily extendable framework for modeling the response at the molecular level together with a simple and robust method to analyze the experimental data. Applied together, these amount to a powerful prescription to probe the effect of inhibitors on signaling cascades.

Our study is multi-disciplinary by its nature, bridging cell biology, quantitative biology, and pharmacology. Therefore, to make our work as accessible as possible, we have suppressed the mathematical details from the main body of the text; instead, they are laid out in the Supplementary Material. The rest of this is organized as follows: we begin by studying a simpler signaling cascade, the JAK-STAT pathway, to demonstrate that cell-to-cell variability correlates with variable response to inhibition. Once we acknowledge the importance of cell-to-cell variability when studying inhibition, we then shift our attention to the more complex MAP kinase cascade, where we explicitly demonstrate the existence of two qualitatively different modes of inhibition. To understand the origin of the different modes of inhibition, we develop a coarsegrained model of CD8^+^ T-lymphocyte activation and inhibition and compare it to our data. We conclude the results section by demonstrating that the different modes of inhibition of the signaling network map to different functional fates, as measured by a simple cell proliferation assay. We thus establish the link between inhibiting a heterogeneous cell population, the targeted signaling network structure and role of the specific inhibited enzyme in it, and the functional consequences of such inhibition.

## Results

### Endogenous variability of STAT5 creates variability in phospho-STAT5 response to JAK inhibition

A reductionist approach posits that the properties of a signal transduction pathway in living cells should be deductible from the biochemistry of its components working *in concert*. However, traditional methods such as *in vitro* assays of enzyme extracts and ensemble average measurements (*e.g*. western blot) do not incorporate the inherent biological complexity of cells or the required resolution, and therefore fall short of a detailed biochemical characterization of chemical inhibitors. To illustrate this issue, we investigated the biochemistry of JAK-induced STAT5 phosphorylation in individual T lymphocytes stimulated with the cytokine Interleukin 2 (IL-2, Fig. 1A). We focus on this pathway for three reasons: (i) its biological function is important, corresponding to anti-apoptotic and proliferative signals [30]; (ii) its clinical relevance in inflammatory diseases [2, 3] and cancer [31]; (iii) its the molecular components are well documented [32].

**Figure 1:**
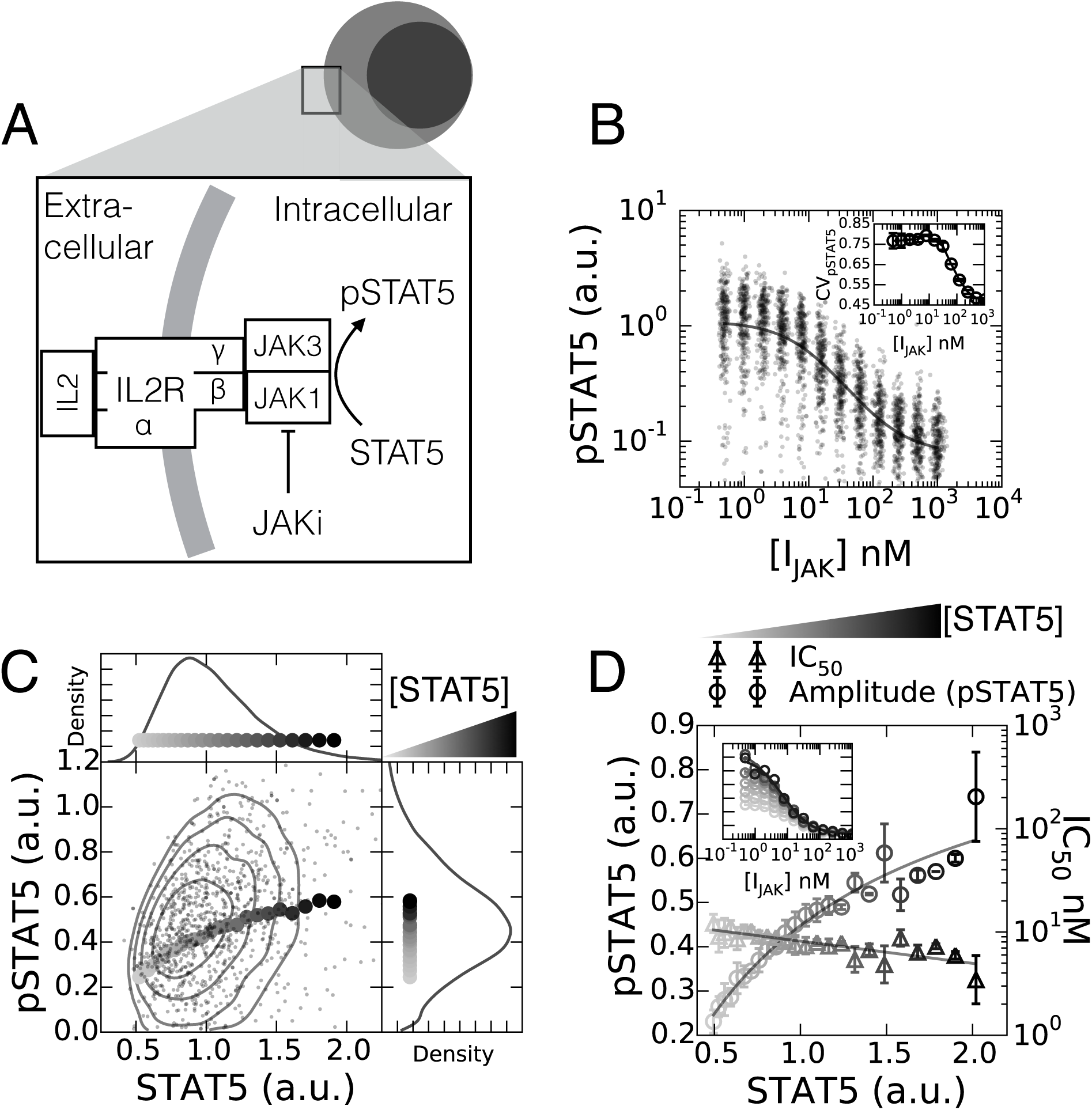
Variability of endogenous protein abundance correlates with single cell response to chemical inhibition. (A) IL-2 stimulation of the JAK-STAT pathway. (B) Single cell pSTAT5 abundance in response to JAK inhibitor AZD1480. Inset, the coefficient of variation (CV) response to inhibition. (C) Single cell contour plot of total STAT5 abundance and pSTAT5 in cells not treated with inhibitor, [I_jak_] = 0. Curve shows the resulting geometric mean of the pSTAT5 abundance conditioned on STAT5 abundance per cell. (D) Cell-to-Cell Variability Analysis reveals that the pSTAT5 response amplitude is correlated with STAT5 abundance. In addition, the sensitivity of cells to inhibition (IC_50_) exhibits a small negative correlation with STAT5 abundance (errorbars are standard deviation of experimental duplicates).

In order to monitor the JAK/STAT signaling response to JAK inhibition - we prepared *ex vivo* mouse primary T cell blasts and exposed them to saturating amounts of the cytokine IL-2 (2 nM), followed by two-fold serial dilutions of AZD1480 (I_jak_). We found that the average response follows an inhibitory hill function with an estimated half effective inhibition concentration (IC_50_) of 8.2 ± 0.5 nM (Fig. 1B). In this preliminary characterization we assumed that the hill coefficient is exactly one. A hill coefficient of one indicates that the inhibition of STAT5 phosphorylation can be described by the drug simultaneously binding and deactivating the kinase.

The phospho-STAT5 (pSTAT5) response of individual cells to JAK inhibition decreases smoothly and unimodally with increasing doses of drug (Fig. 1B). We characterized the variability of cell responses by the coefficient of variation (CV), a measure of the standard deviation with respect to the mean pSTAT5 response. In the absence of drug the CV is 0.77 ± 0.004, and depreciates with increasing doses of inhibitor (Fig. 1B inset). The concomitant decrease in the mean response and CV_pstat5_ contradicts the stochastic properties of chemical reactions. Indeed, diversity in the abundance of pSTAT5 originating from physicochemical mechanisms is expected to exhibit Poisson statistics, meaning that the CV_pSTAT5_ should behave as the inverse square root of the mean [33, 34]. Therefore, in contrast to our observations, if the origin of the noise were Poissonian, CV_pSTAT5_ originating from these simple Poisson properties would increase, rather than decrease, with increasing inhibitor dosage. Consequently, we conclude that individual clones generate diverse levels of pSTAT5 from biological sources of diversity, *i.e*. protein variability, as opposed to the intrinsic stochasticity of chemical reactions.

Next we asked whether the variable abundance of STAT5 can explain pSTAT5 variability in response to JAK inhibition. We simultaneously monitored both STAT5 and pSTAT5 in individual cells, and measured the average pSTAT5 abundance in subpopulations of cells with similar STAT5 abundances, a technique referred to as Cell-to-Cell Variability Analysis (CCVA) [28, 29]. We found that the geometric mean of pSTAT5 correlates with STAT5 abundance in the absence of JAK inhibitor (Fig. 1C). We then investigated how varying abundances of STAT5 influence both the JAK inhibitor dose response amplitude and the half effective inhibitor concentration (IC_50_). We found that the amplitude of pSTAT5 response increased with STAT5 expression, while the IC_50_ reduced exponentially with a scale of approximately -2.0 (STAT5 a.u., Fig. 1D). Hence, by monitoring the extent of drug inhibition at the single cell level, we establish new experimental observations regarding signal inhibition.

CCVA established the dependence of pSTAT5 on the endogenous (variable) STAT5 abundance. We leveraged this observation to develop a biochemical model of inhibition in live cells. Specifically, we tested three simple biochemical models that may account for the transmission of STAT5 variability to pSTAT5 levels per cell, and the biochemical mechanism of JAK inhibition by AZD1480 (Fig. 2A, see Supplementary Material Sections 1.1-1.3 for derivations). Each mechanism represents unique interactions between the JAK, STAT5, and I_jak_ - the noncompetitive inhibitor binds to JAK independent to the presence of STAT5, the uncompetitive inhibitor only binds to the JAK-STAT5 complex, and the competitive inhibitor binds to JAK which prevents STAT5 from binding (Fig. 2A). We found that a noncompetitive inhibition model for AZD1480 action best described our experimental observations (Fig. 2B). This observation is supported by the fact that AZD1480 acts by competing with ATP for occupancy of the ATP binding pocket of JAK, and does not compete with STAT5 [35]. Furthermore, it was necessary to account for the physiological variability in STAT5 substrate availability to account for the cell-to-cell variability in pSTAT5 inhibition (Fig. 2C,D). Lastly, we validated that our model could account for the small dependence of the IC_50_ on STAT5 expression. We found agreement between our IC_50_ measurements in Fig. 1D with the estimated IC_50_ from our model (Fig. 2E).

**Figure 2:**
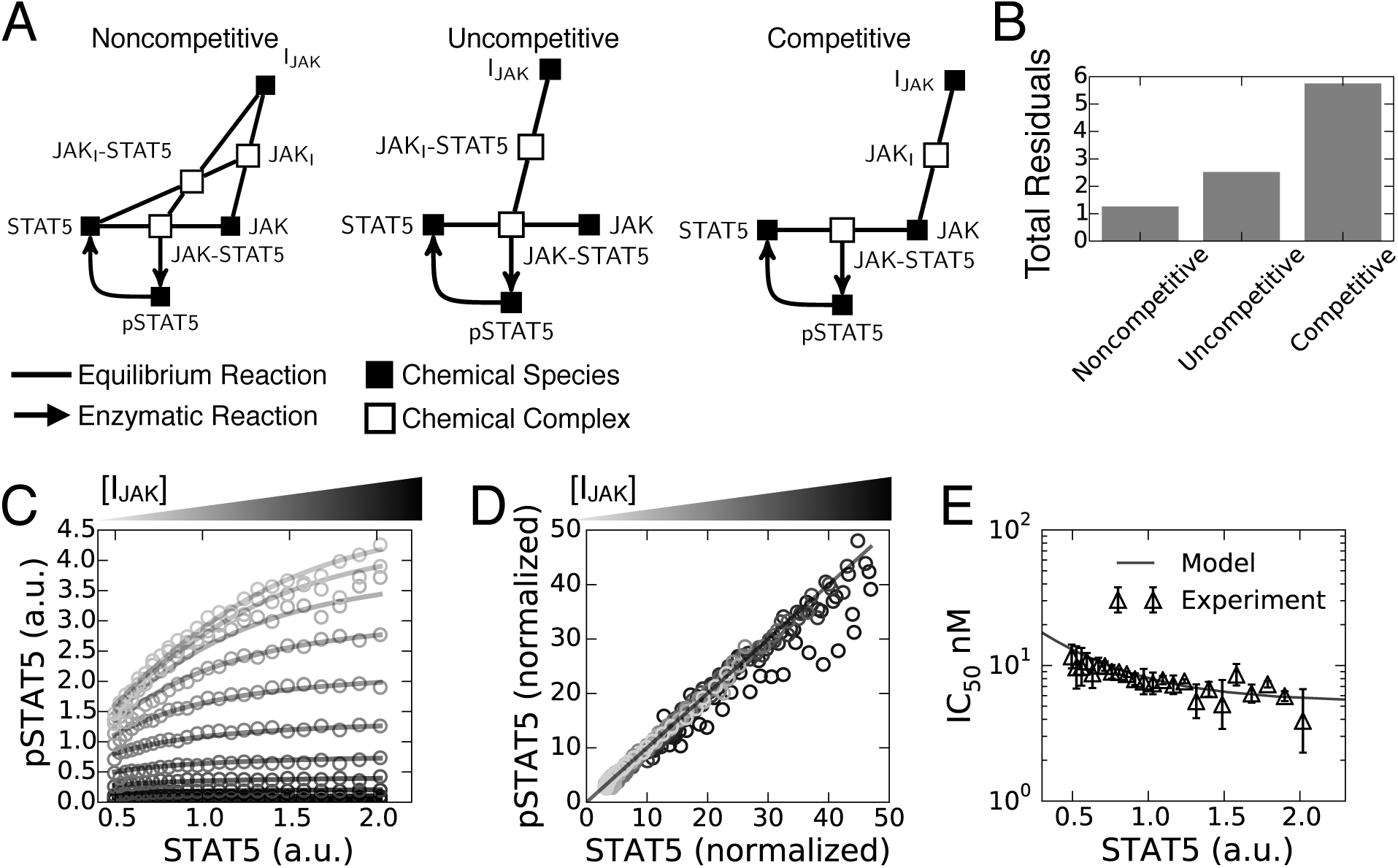
CCVA reveals the most likely mechanism of AZD1480 in live single cells. (A) Model diagrams that represent three possible mechanisms of inhibition. (B) Each model was tested against our data by measuring the sum of squared residuals (Total Residuals) between our model predictions and the data points - a lower value means better agreement between model and data. The model was fit to all the data point presented in (C). (C) Overlay of data (circles) with the optimal model and parameter set from fit (lines). (D) A linear transform of the data was derived from the optimal model and the corresponding parameters reveal agreement between model (line) and the data (open circles), (See Eq. S8 in Supplementary Materials for details regarding the nonlinear transformation). (E) Overlay of measured IC_50_ with respect to STAT5 abundance as measured from CCVA analysis of data (triangles; errorbars standard deviation experimental duplicates) and predicted by our optimal model (line).

To summarize, in this section we demonstrated how CCVA parses single-cell phospho-profiling data to validate models of drug inhibition. We employed CCVA here on the JAK-STAT pathway, and found an optimal model of non-competitive binding of inhibitor to kinase, as supported by the literature.

### Single cell measurements reveal diverse modes of inhibition

We proceed by investigating inhibition of a more complex signaling cascade, namely antigen-driven MAP kinase activation in primary T cells. Upon exposure with activating ligands (*e.g*. complex of a peptide with Major Histocompatibility Complex, peptide-MHC presented on the surface of antigen presenting cells), T cells activate their receptors through activation of a SRC Family kinase (Lck). This then triggers a cascade of kinase activation leading to ERK phosphorylation. We chose this model system because its complex network topology and its functional relevance: aberrant activation in the ERK pathway is often involved with oncogenesis [4], making it a key pathway to be targeted with drug inhibitors in multiple tumor settings.

It is possible to decompose T cell receptor mediated ERK signaling into two smaller subnetworks: (i) a receptor proximal signaling cascade with positive and negative feedback regulation, and (ii) the unidirectional MAP Kinase (MAPK) cascade. We now demonstrate that inhibiting enzymes specific to each signaling sub-network produces a unique response in terms of ERK phosphorylation. To show this, we subjected activated T lymphocytes to inhibitors targeting the two signaling sub-networks separately: a SRC inhibitor (Dasatinib) for the receptor proximal component, and a MEK inhibitor (PD325901) for the MAPK component (see Fig. 3A). Importantly, the population-mean response of the cells to each inhibitor resulted in amplitude reduction and a trivial inhibition model (see Fig. 3D,E insets). However, going down to the single-cell resolution, the ppERK response to SRC inhibition (Dasatinib) resulted in an all-or-nothing response (“digital” inhibition, Fig. 3B,D). Conversely, application of a MEK inhibitor (PD325901) resulted in graded responses (“analog” inhibition, Fig. 3C,E) [36]. SRC and MEK inhibitors exhibit markedly different modes of inhibition which do not rely on the exact chemical identity of the administered inhibitor but rather its role in the signaling cascade (see Supplementary Material).

**Figure 3:**
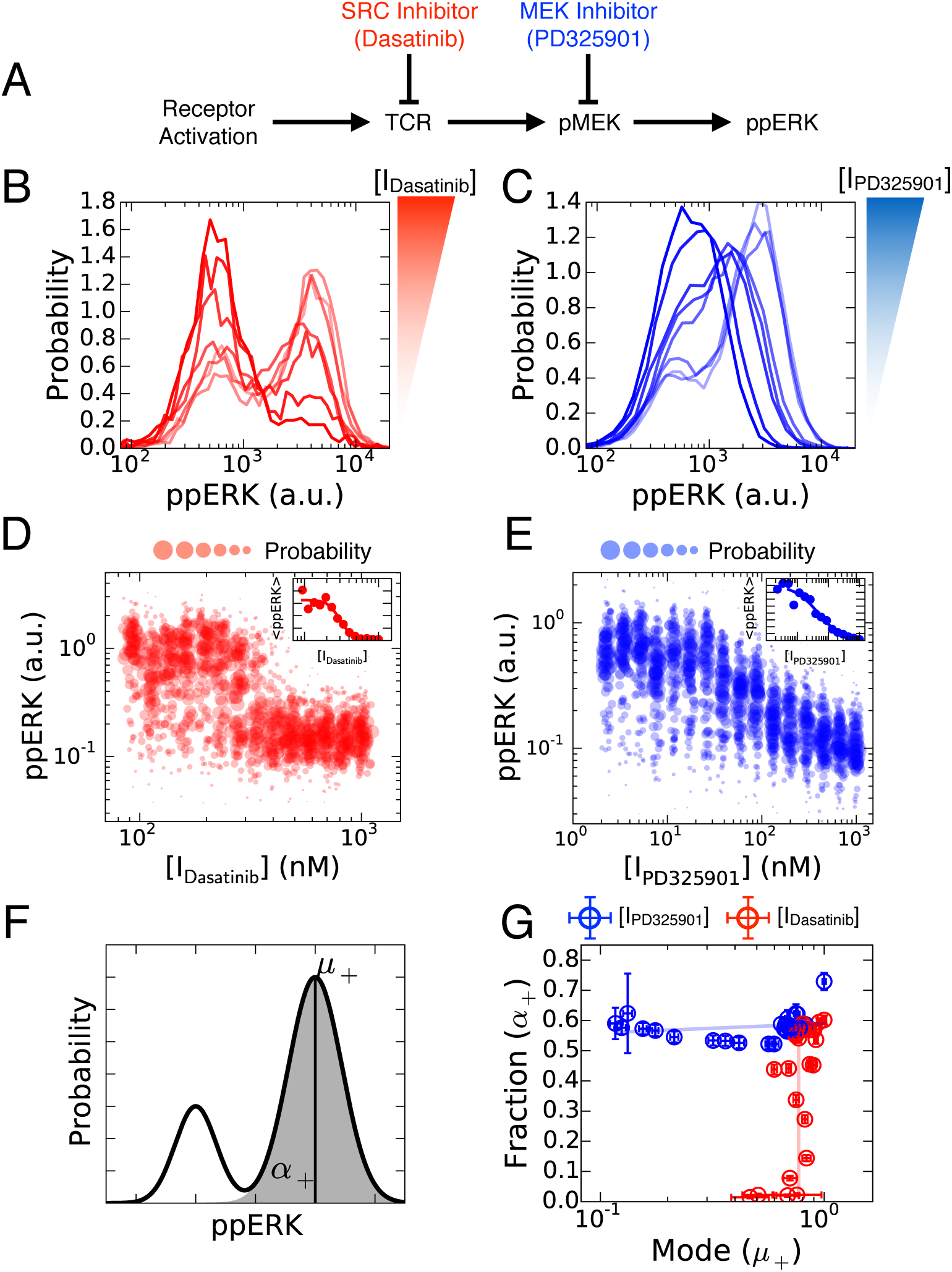
Specific modes of inhibition. (A) The implicit model of inhibitor action. (B) Histograms of single cell response to TCR stimulation and SRC inhibition with Dasatinib. (C) Histograms of single cell response to TCR stimulation and MEK inhibition with PD325901. (D) Single cell dose response of Dasatinib the inset represents the mean response of all cells to a single dose of inhibitor. (E) Single cell dose response of PD325901 - the inset represent the mean response of all cells to a single dose of inhibitor. (F) Single cell data were modeled as a mixture of Gaussian distributions. *α*_+_ parameterizes the fraction of activated cells, while *μ*_+_ parameterizes the average abundance of ppERK among activated cells. (G) The (*μ*_+_,*α*_+_) plane shows the orthogonal modes of inhibition (errorbars are standard error of mean from 100 samples of 500 T cells per dose of inhibitor chosen randomly and with replacement).

We characterized the two modes of inhibition by fitting the distribution of ppERK amount per cell to a mixture of two Gaussians. The relevant statistics can be summarized by two parameters - *α*_+_ which represents the fraction of activated cells (Fig. 3F) and *μ*_+_ representing the mean ppERK levels among activated cells. We carried out this analysis for each dose of inhibitor. In Fig. 3G, we report that the MEK inhibitor operates solely upon the mean, *μ*_+_, of ppERK abundance among activated cells, which we define as analog inhibition of ERK activation. In contrast, the SRC inhibitor operates solely upon the fraction of active cells, *α*_+_, which we define as digital inhibition. To summarize, by utilizing single-cell measurements, we were able to demonstrate that there exist two modes of inhibition in the MAPK signaling cascade, digital and analog, each of which are associated with the sub-network the targeted kinase belongs. Each mode of inhibition can be associated with the unique inhibition of proximal and distal kinases respectively within the ERK cascade. We proceed to examine if this effect can be captured by the properties of the respective sub-networks and whether it maps to a functional output.

### Sub-network context of the targeted enzyme determines response to inhibition

We explored whether the two distinct modes of inhibition observed in Fig. 3 originate in the context of the targeted enzymes, *i.e*. by the position of the enzyme undergoing inhibition within the transduction cascade. For this, we developed a coarse grained model which accounts for ERK phosphorylation downstream of SRC activation [24, 37]. Our model explicitly incorporates measurable quantities (*e.g*. abundance of CD8), control parameters (e.g. antigen concentration), and the inhibitor targeted species [38], while incorporating uncontrollable or unmeasurable quantities into the phenomenological species “activated SRC”. The graphical representation of our model (Fig. 4A) emphasizes the two sub-networks acting here. First, SRC* is controlled by competing positive and negative feedbacks that are abstractions of negative feedback of active SHP-1 phosphatase [24] and positive feedback associated with immune receptor signaling [37, 39]. Whereas, MEK and ERK are activated upon formation of SRC* in an unidirectional manner, without feedback (see Supplementary Material Section 3.3 for experimental evidence for our uni-directional MAPK signaling assumption.). These modeling components encompass key features of ERK activation in the context of antigen activation in T lymphocytes.

**Figure 4:**
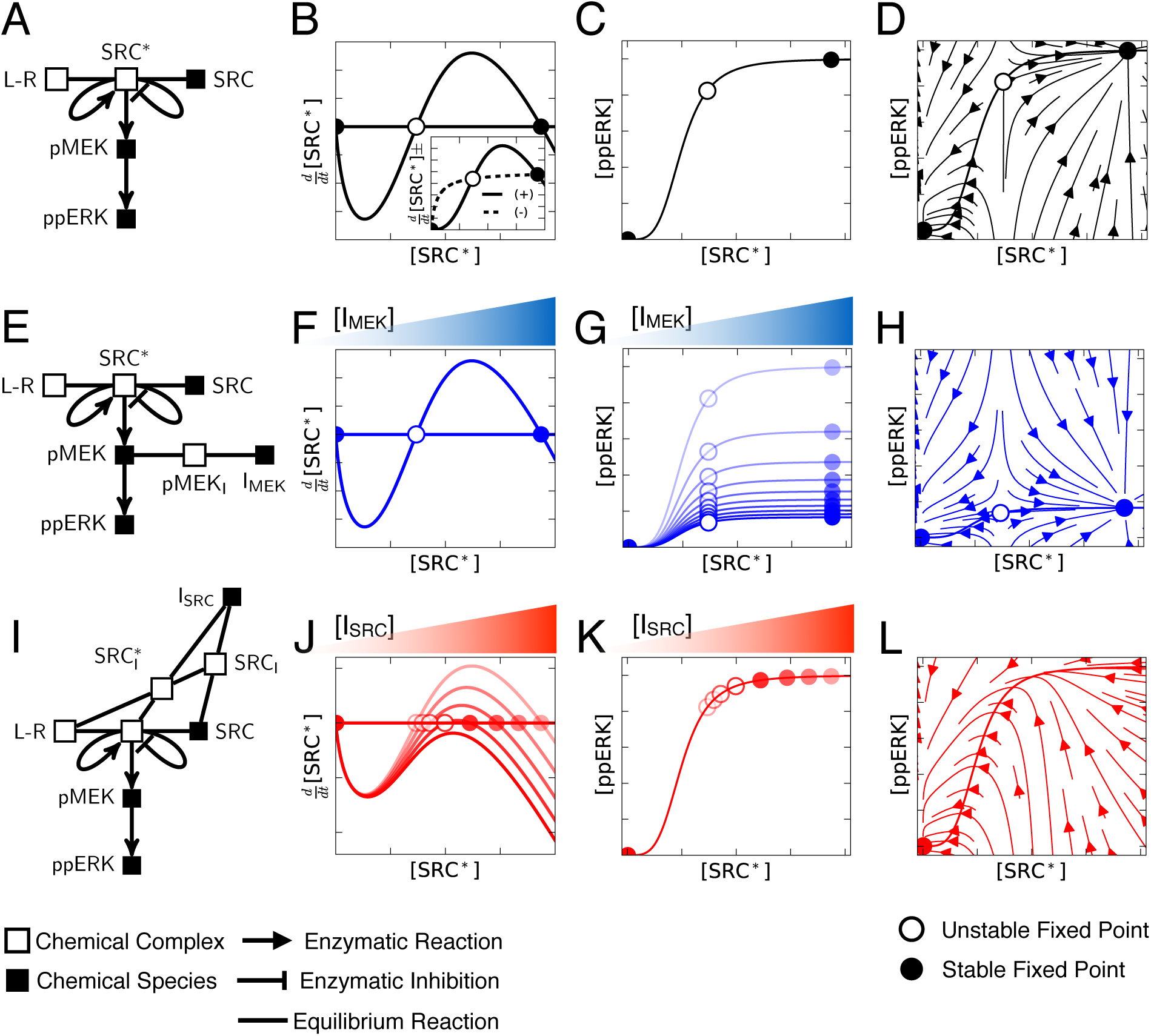
Sub-network context of the targeted enzyme determines response to inhibition. (A) Model diagram of signaling network. (B) Phase plane of SRC* with respect to zero flux (horizonal line). The inset shows the behavior of both the positive and negative model fluxes. (C) Functional response of ppERK to changes in SRC* abundance. (D) Instantaneous reaction velocities given ordered pairs of (SRC*, ppERK) shows the dynamic behavior of the model system. (E) Model diagram for MEK inhibition of signaling. (F,G) Analogous to representations of model behavior as (B,C) but for different doses of MEK inhibitor. (H) Instantaneous reaction velocities for maximal MEK inhibition. (I) Model diagram for SRC inhibition. (J,K) Analogous representations of SRC inhibition as (F,G) for MEK inhibition. (L) Instantaneous reaction velocities for maximal SRC inhibition show that the dynamics support a single fixed point at 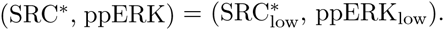.

To assess the properties of our competing feedback model we constructed phase diagrams of active SRC that demonstrate how varied quantities of active SRC map to ERK phosphorylation. Active SRC accumulates upon engagement of the activating ligand with its kinase-bearing receptor. This subsequently activates both positive and negative feedbacks driving further accumulation or extinction of SRC*. The dynamics of accumulation of SRC* can be summarized in a phase diagram (Fig. 4B), that illustrates the influence of both feedbacks (for details and the derivation of these phase diagrams see the Supplementary Material Section 2). The model parameters are set so that the negative flux (*i.e*. the change in time of SRC* levels) rises and saturates at lower levels of SRC* than the positive flux. This staggering of the positive and negative fluxes as a function of SRC* causes them be equal at three points in the phase diagram, *i.e*. there are three “steady states” or fixed points in our model. By plotting the net flux as a function of the active complex SRC*, we assessed the stability of the fixed points. The dynamics are such that SRC* always converges to the extreme points 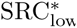 and 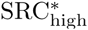 (stable fixed points), while diverging from the center point 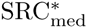 (unstable fixed point). Hence, our coarse-grained model encapsulates the bistability in SRC* formation.

We model ERK activation by assuming that the active complex SRC^*^ triggers the enzymatic phosphorylation of MEK, which then phosphorylates ERK (Fig. 4C). In Fig. 4D, we represent the dynamic trajectory of this signaling pathway for varied initial conditions: such a flow diagram illustrates the stability of the low and high states in the (SRC*,ppERK) plane and the instability of the intermediate point. Overall, our coarse grained model of ERK activation upon ligand engagement generates two stable fixed points corresponding to either low or maximum ppERK, consistent with our experimental results.

Next, we tested whether our coarse-grained model can predict ppERK response to drug inhibition. Application of the MEK inhibitor to our model (Fig. 4E) supports our experimental observations, as MEK inhibition does not influence the bistability of the activated kinases SRC* (Fig. 4F). Increasing the MEK inhibitor dose shows continuous reduction in the amplitude of ppERK response (Fig. 4G), without affecting the bistability in ppERK. The dynamic properties supporting the bistability in ppERK are preserved in the presence of the MEK inhibitor (Fig. 4H). Our model is validated with the experimental observation in that the MEK inhibitor only reduces the mean quantity of ppERK over the population of activated cells, *i.e*. it inhibits the ERK pathway in an analog manner.

Our model highlights that SRC is the kinase crucial for the bistability of the active complex SRC^*^, resulting in a signaling context fundamentally distinct to that of MEK. Inhibition of SRC reduces the positive flux which generates SRC*(Fig. 4I), and consequently reduces SRC* at the high fixed point. We find that increasing the dose of the SRC inhibitor decreases SRC^*^ until, at a critical dose, the high fixed point and the unstable fixed point annihilate one another (Fig. 4J). Therefore, a dose of SRC inhibitor greater than the critical dose leaves the system with only a single fixed point, 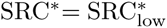 (Fig. 4J). Interestingly, despite the continuous reduction of the 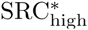 stable fixed point with increased dosage of SRC inhibitor, the quantity of ppERK remains essentially unchanged until the inhibitor is greater than the critical dose (Fig. 4K). For doses of SRC inhibitor beyond the critical dose the signaling network only supports a single low quantity ppERK (Fig. 4L). Hence, SRC inhibition results in a binary output that is identical to that observed in the data: our model is consistent with the digital nature of Dasatinib as a SRC inhibitor.

Our model assumes that interactions of molecular inhibitors with their target enzyme all act as noncompetitive inhibitors (consistent with *in vitro* characterization of these small molecules). Yet, despite these locally identical mechanisms of inhibition, the model successfully accounts for the two distinct modes of inhibition of ERK in our experimental findings. Thus, taken together, the experimental results and theoretical model demonstrate that enzymatic context is essential to understand and parameterize inhibitor function.

### Variability in protein abundance diversifies sensitivity of signaling pathway to targeted inhibition

Using our coarse grained model, we sought to explore how the endogenous variability of SRC abundance would diversify the response of individual cells to inhibition. Our model predicts that the effective quantity of SRC determines whether the (SRC, ppERK) phase diagram has a single or three fixed points - as a result it represents a bifurcation parameter. By analogy, endogenous variation of SRC positions cells either above or below the critical threshold of SRC required for bistable signaling (Fig. 5A). We tested this hypothesis by correlating CD8 and ppERK of activated T lymphocytes. In T lymphocytes, Lck - a SRC family kinase, is recruited together with CD8 to trigger response to antigen, therefore we treat CD8 abundance as a proxy for the effective abundance of SRC in individual cells. Indeed, measuring CD8 for a single dose of SRC inhibitor shows that cells with elevated quantities of CD8 are more likely to have ppERK signal (Fig. 5B), a result that is consistent with previous experimental and theoretical observations [38].

**Figure 5:**
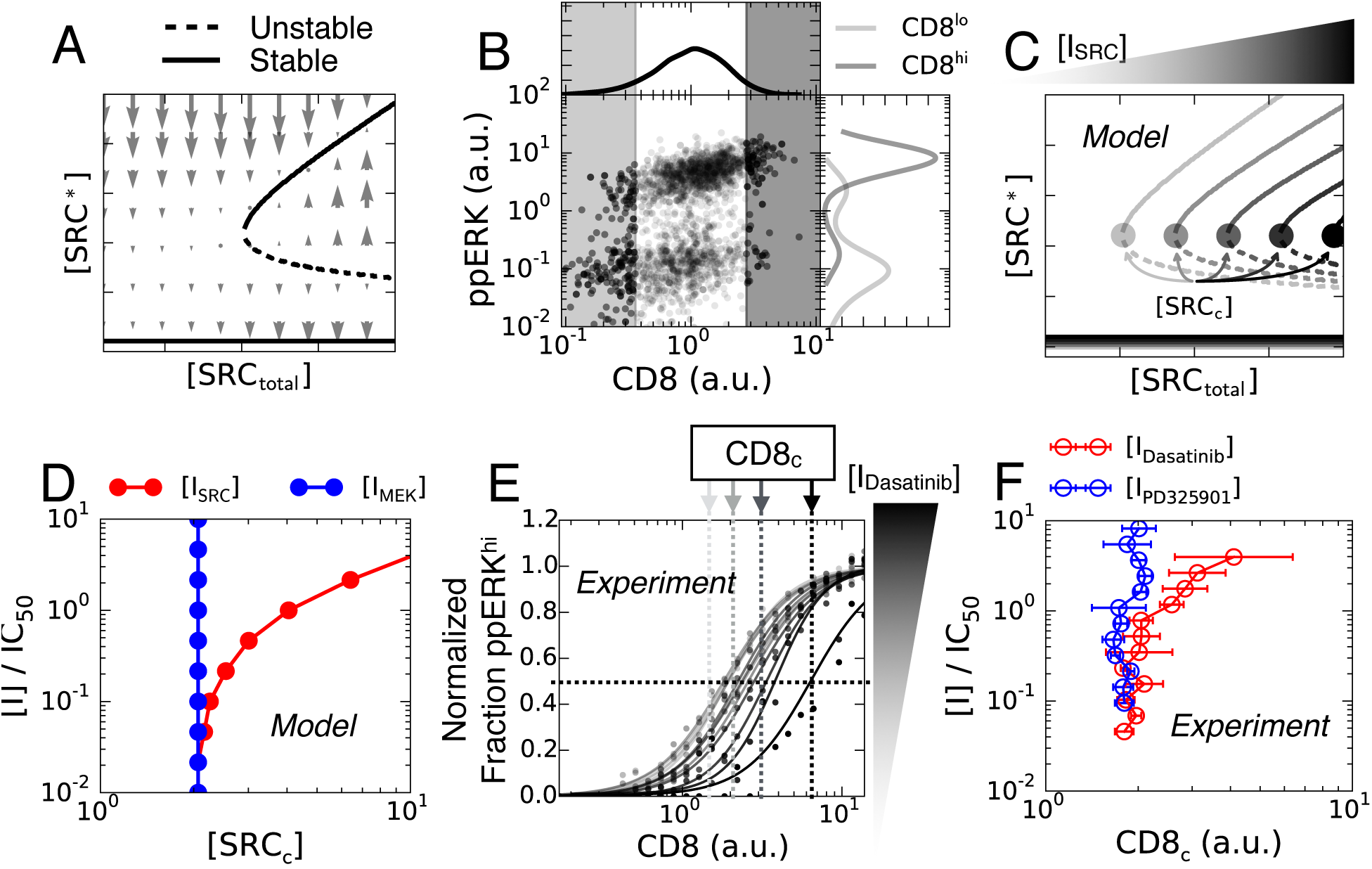
Protein variability tunes sensitivity of cells to inhibition. (A) The abundance of SRC_to_tai controls the number of fixed points of the model, and consequently is a bifurcation parameter. (B) Flow cytometry measurements of T cells concomitantly labeled for CD8 (a proxy SRC) and ppERK (pT202, pY204) shows signaling dependence to endogenous expression of CD8. (C) Model predictions of SRC_c_ dependence on SRC inhibitor dose. (D) Model predicts scaling of SRC_c_ with increased SRC inhibitor doses. Intuitively, MEK inhibitor does not influence SRC_c_. (E) CCVA of T cells concomitantly labeled for CD8 and ppERK (pT202, py204) treated with various doses of the SRC inhibitor Dasatinib. (F) Quantification of the half effective abundance of CD8 (CD8_c_) required for ppERK activation in T cells treated with either MEK inhibitor (PD325901) or SRC inhibitor (Dasatinib). Errorbars quantify one standard deviation from the mean of experimental duplicate measurements.

Extending this observation, our model suggests an interesting possibility: that variability of CD8 expression in single cells is sufficient to generate disparate sensitivities to drug inhibition. The bifurcation diagram for each drug dose, Fig. 5C, shows that the minimum quantity of SRC sufficient for the bistability, SRC_c_, increases with increasing drug dose. Consequently, a cell with a higher abundance of SRC will be more tolerant to inhibition because of simple dosing of the effective abundance of available SRC (Fig. 5D) - requiring higher inhibitor dosage to experience any reduction in signaling. We confirmed this qualitative prediction by correlating the critical amount of CD8, labeled CD8_c_, with drug dose; the MEK inhibitor reduced ERK activation independently of the abundance of CD8 whereas higher concentrations of SRC inhibitor were required to inhibit (Figs. 5E,F).

### Diverse cell signaling responses to inhibition translates to diverse proliferative responses to inhibition

Having established the existence of distinct modes of inhibition of the ERK pathway, we conclude the results section of this *Communication* by posing an important challenge to our finding: do these distinct modes of inhibition entail a functional ramification? Upon phosphorylation, ppERK migrates from the cytosol to the cell nucleus where it induces the expression of the Immediate and Early Genes (IEGs, *e.g*. cFOS). IEGs constitute a set of genes that facilitate cell cycle entry and cell division [40]. Hence it is reasonable to expect that inhibiting the ERK pathway will impact cell proliferation. But will cell proliferation, which happens on the scale of days, be sensitive to the different modes of inhibition, which happen on the scale of minutes ? Our signaling results (Fig. 3) suggest the following hypothesis: that MEK inhibition, which produces intermediate levels of ppERK, will slow down induction of IEGs, and as a result would increase the time to cell division. In contrast, SRC inhibition, which reduces the fraction of cells getting activated, will reduce the number of cells entering cell cycle, without affecting the overall cell division in activated cells (cf Fig. 3).

We tested this hypothesis by quantifying the proliferation of T cells after 48 hours of *in vitro* culture under concomitant antigen stimulation and drug exposure. We used flow cytometry to monitor cell activation and division by measuring cell size (FCS-A), the levels of the CD8 co-receptor on the surface of cells (proportional to fluorescence intensity), and the fluorescence of T cells that were tagged before activation with an amine-reactive fluorescent dye (CTV or CFSE, the dye gets diluted by 2 fold at each cell division). Upon activation, T cells increase both their size and CD8 expression, providing a clear criterion (Fig. 6A) that separates inactive and active cell populations, whose numbers can be quantified as N^-^ and N^+^, respectively. Among the active fraction of cells we analyze the number of cells 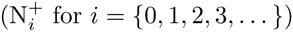 undergoing *i* divisions as measured by CTV or CFSE dilution (Fig. 6B). By computing both the mean number of divisions and the fraction of activated cells for each dose of each drug, we could plot the two hypothesized modes of long timescale inhibition.

**Figure 6:**
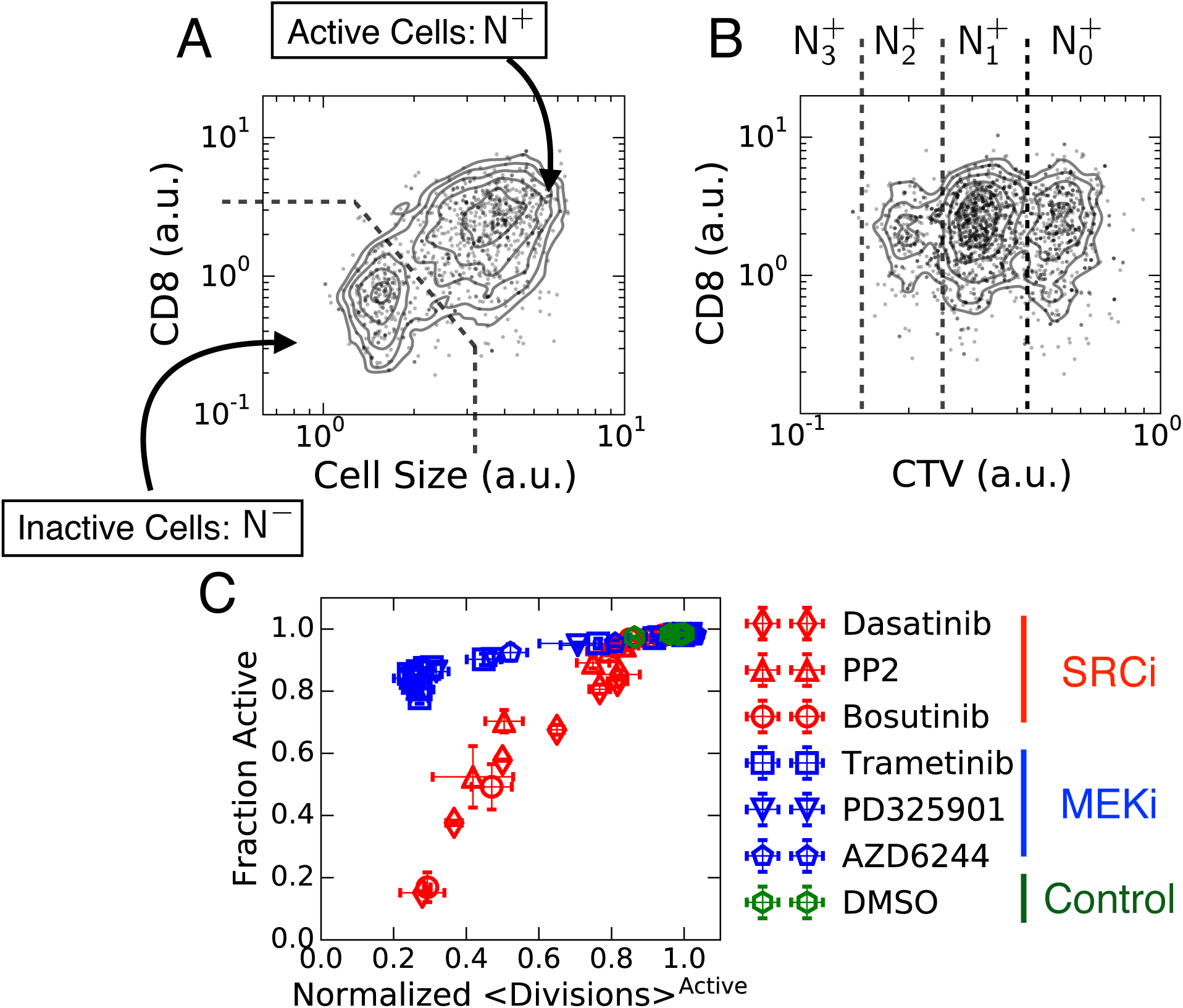
Short timescale modes of inhibition translate to long timescale proliferative response. (A) Identification of activated T cells after 48 hours of concomitant treatment with antigen and the respective inhibitor measured by flow cytometry. Upon activation, cells increase size, measured by forward scatter area (FSC-A), and upregulation of CD8. (B) CellTrace Violet (CTV) dilution identifies active cells that have divided *n* times. (C) Quantification of fraction of active cells and mean number of divisions, among activated cells, for each inhibitor (See Supplementary Material Section 4 for derivation). Errorbars quantify one standard deviation from the mean of experimental duplicate measurements.

Representing the data as fraction activated vs. mean divisions, demonstrates that the disparate modes of inhibition for signal transduction map to the proliferative timescale (Fig. 6C). To be more explicit, we found that dosing of MEK inhibitor reduces the average number of divisions among activated cells, while the dominant feature of the SRC inhibition is the distinct reduction of the number of activated cells. This is not the exclusive feature observed in our data, since intermediate doses of SRC inhibitor do also reduce the mean divisions (possibly because of the unaccounted signaling transduction pathways dependent on TCR activation, *e.g*. PI3K and AKT [41]). Crucially, application of MEK and SRC inhibitors shows grouping of the proliferation data when represented as fraction activated vs. mean division number. We then found these results to be a general property of MEK and SRC inhibitors in our system by including the following: Bosutinib, PD325901, PP2, Trametinib and AZD6244; most these drugs are either presently clinically used or in various stages of clinical trials. Indeed, Fig. 6C shows an astonishing degree of agreement in-between the SRC inhibitors, in-between the MEK inhibitors, and at the same time, a very clear divergent behavior of the two families. We conclude that our measurements support the hypothesis that the impact of MEK/SRC inhibition on cell proliferation recapitulate the two modes of inhibition that we documented with short-term signaling response in Fig. 3.

## Discussion

In this study we combine theoretical and experimental approaches to probe mechanisms of inhibition in signal transduction. We used single cell phospho-profiling and Cell-to-Cell Variability Analysis (CCVA) [28] to characterize the biochemical details of small molecule chemical inhibitors within living cells; such detail was so far limited to *in vitro* enzymatic assays. We uncovered a generic mechanism in which targeted enzyme inhibition manifests in two distinct patterns of inhibition, which we label “digital” vs. “analog”. Lastly, we probed the biological significance of these results by correlating short timescale signaling behavior with unique modes of inhibition of cellular proliferation.

Using single cell phospho-profiling and CCVA we were able to perform detailed and mechanistic characterization of cellular responses to targeted inhibition in primary cells. Specifically, we show how to utilize CCVA to mechanistically characterize the biochemical interaction between the enzyme target and the inhibitor. We confirmed that AZD1480 is a potent non-competitive, with respect to STAT5, inhibitor of JAK-STAT signaling in IL-2 stimulated primary T lymphocytes (Fig. 2) and that pSTAT5 levels and drug efficacy depend on varied levels of endogenous STAT5. In addition, we demonstrated how the organization of reactions in biochemical networks in a more complex signaling cascade can determine markedly different cellular responses to inhibition (Figs. 3,4).

Although Albeck *et al*. also found that inhibition of different enzymes manifest to digital or analog signaling responses [26], we propose a mechanistic model which attributes these disparate responses to the context of the targeted enzyme. Essentially, the overall network response to inhibition is determined by the dynamic properties associated with the targeted enzymes’ location in the larger biochemical network (Fig. 4). Furthermore, we show how our short timescale signaling behavior translates to novel long timescale proliferative response to inhibition (Fig. 6). As a result, our method extends the characterization of inhibitors from the current state-of-the-art *in vitro* assays to primary single cells, and sheds light on the nonlinear signaling responses of the biochemical network structure being perturbed.

Building upon our initial findings, we extended our mechanistic models and used CCVA to demonstrate how cells utilize the endogenous variability of protein abundance to generate disparate responses to singular perturbations. In context to inhibition, we found that variation in enzyme substrate (STAT5) abundance established diverse signaling amplitudes and varied the sensitivity of single cells to inhibition (Figs. 1,2). We then extended our mechanistic model of SRC inhibition and found that the variability of SRC expression operates on cells as a bifurcation parameter, which controls the number of possible steady states of the signaling network. As a result, cells that had elevated abundance of SRC were more tolerant to inhibition (Fig. 5). The extent in which these mechanisms of diversity provide resilience of populations to inhibition at longer timescales remains an open question. However, our findings are of practical importance: there are numerous examples of biological systems that utilize protein abundance to generate phenotypic variability, as noted in *e.g*. [42, 43, 44, 45, 46]. Similarly, there exist abundant single cell observations showing heterogeneous responses to inhibition [16, 21, 22].

Our method facilitates the extension of *in vitro* kinase assays to cellular systems and motivates a transition from phenomenological characterization of drug response at the single cell level, into mechanistic and functional understanding. By using our combined approach of CCVA and development of mechanistic models to characterize drugs in primary cells, we were able to unravel fundamental chemical and biological processes. In particular, our method is especially useful when probing the functional consequences (on long timescales) of molecular perturbations (as experienced by cells on short timescales). When applied, we successfully showed how the SRC and MEK inhibitors cluster on two distinct curves, which are easy to interpret as distinct modes of inhibition. Since it is unlikely that the ERK pathway is the only cellular pathway exhibiting distinctly different modes, we expect that our method will prove useful in characterizing other inhibitor-pathway combinations, hopefully teasing out more novel modes of inhibition.

## Methods and Methods

### Mice and Cells

Primary splenocytes and lymphocytes were harvested from C57BL/6N (B6; Taconic Farms), B10A wild type (B10A; Taconic Farms), OT-1 TCR transgenic RAG2^-/-^ (Taconic Farms), and 5C.C7 TCR transgenic RAG2^-/-^ (Taconic Farms) mice and cultured up to 10 days. Mice were bred, maintained, and euthanized at Memorial Sloan Kettering Cancer Center (MSKCC) in compliance with our animal protocol. The animal protocol was reviewed and approved by the Institutional Animal Care and Use Committee (IACUC) of the Memorial Sloan Kettering Cancer Center (New York NY). The protocol number is 05-12-031 (last renewal data: December 23rd 2013). RMA-S TAP-deficient T cell lymphoma cell line was used as antigen presenting cells for signaling experiments [24].

### Antibodies and Cell Stains

Cells were labeled with primary antibodies against doubly phosphorylated ERK 1/2 (p-T202, p-Y204; clone E10), phosphorylated MEK 1/2 (p-S221; clone 166F8), or phosphorylated STAT5 (p-Y694; clone C11C5) - purchased from Cell Signaling Technology (Beverly, Massachusetts). The primary antibody against STAT5 was purchased from Santa Cruz Biotechnology (Santa Cruz, California). Secondary antibodies tagged with fluorescent molecules were purchased from Jackson ImmunoResearch (West Grove, Pennsylvania). Surface markers CD8*α* (clone 53-6.7) and CD4 (clone RM4-5) tagged to fluorescent molecules were purchased from Tonbo biosciences (San Diego, California). Cell proliferation was measured by dilution of either CellTrace^TM^ Violet (CTV) or Carboxyfluorescein *N*-succinimidyl ester (CFSE) proliferation kits purchased from Molecular Probes. Cell viability was assessed with Live/Dead^®^ Near-IR kit purchased from Molecular Probes.

### Small Molecule Chemical Inhibitors

The SRC inhibitors PP2 and Bosutinib as well as the MEK inhibitor PD0325901 (PD325901) were purchased from Sigma-Aldrich. The MEK inhibitors Trametinib and AZD6244 were generous gifts from Neal Rosen (MSKCC). The JAK inhibitor AZD1480 and SRC inhibitor Dasatinib were purchased from Selleckchem.

### Additional Reagents

Supplemented RPMI-1640 media was prepared by MSKCC core media preparation facility and was used for all cell cultures and experiments. Media was supplemented with 10% fetal bovine serum, 10 *μ*g/mL penicillin and streptomycin, 2 mM glutamine, 10 mM HEPES (pH 7.0), 1 mM sodium pyruvate, 0.1 mM non-essential amino acids, and 50 *μ*M *β*-mercaptoethanol. Cell were stimulated with with interleukin 2 (IL-2; eBioscience). TCR activating ligands K5 MCC peptide (K5): ANERADLIAYFKAATKF (T lymphocyte 5C.C7 agonist) and ovalbumin peptide SIINFEKL (T lymphocyte OT-1 agonist) were purchased from GenScript. Cells were chemical fixed and permeabilized following signaling experiments with 2% paraformaldehyde (PFA; Affymetrix) and 90% methanol (MeOH). Cells were stained with antibodies and suspended in FACS buffer for flow cytometry measurements. FACS buffer consists of 10 % fetal bovine serum (MSKCC core media preparation facility) and 0.1% sodium azide in PBS. Ficoll-Paque PLUS (GE) was used to purify live cells in culture.

### Primary cell culture

5C.C7 and OT-1 primary cells were cultured *ex vivo* with peptide pulsed antigen presenting cells (APCs) from irradiated (3,000 RAD) B10A and B6 mice, respectively. APCs were pulsed overnight with 1 *μ*M K5 peptide for 5C.C7 activation and 1 *μ*M SIINFEKL for OT-1 activation prior to irradiation. Cells were purified by Ficoll-Paque gradient centrifugation and given exogenous IL-2 (1 nM) every other day. All cells were cultured at 37° C and 5% CO_2_ in supplemented RPMI and used for experiments within 7 days of activation.

### Single cell inhibition of signal transduction assay

#### JAK inhibition

The pSTAT5 response to JAK inhibition was measured using primary 5C.C7 derived T lymphocytes. Cells were aliquoted in 96 well v-bottom plates with exogenous IL-2 (working dilution 2 nM) for 10 minutes and kept at 37°C. The cells were then treated the JAK inhibitor AZD1480 for 15 minutes at 37°C. Followed by 15 minutes of fixing in 2% PFA on ice. The cells were then permeabilized in 90% MeOH and stored at -20°C until staining for flow cytometry.

#### SRC and MEK Inhibition

The ppERK response to SRC and MEK inhibition was measured using primary OT-1 T lymphocytes activated with RMA-S antigen presenting cells. RMA-S cells were suspended in culture with 1 nM SIINFEKL peptide for 2 hours at 37°C, 5% CO_2_, and on a rotator to guarantee mixing. During this time we labeled OT-1 cells with an amine reactive dye, CTV, according to the manufacture’s protocol (Molecular Probes). This fluorescent tag was used to identify OT-1 cells in silico. We rested the OT-1 cells one hour after CTV staining, and then distributed them in a 96 well v-bottom plate. Each well was given various doses of SRC inhibitor and MEK inhibitor and kept at 37°C for 5 minutes. Following the 5 minute exposure to the inhibitors, we added the peptide pulsed RMA-S (10 RMA-S to 1 OT-1 T cell) and pelleted by centrifugation for 10 seconds at 460 rcf at room temperature. This step guaranteed that both cell types, OT-1 and RMA-S, came into contact. The cells were allowed to activate for 10 minutes, followed by fixing on ice in 2% PFA, and then permeabilized and stored in 90% MeOH at -20°C.

### Proliferation assay

The proliferative response of OT-1 T lymphocytes to SRC and MEK inhibitors was measured by the dilution of the amine reactive dyes CTV or CFSE. Splenocytes from B6 mice were used as antigen presenting cells (APCs). Once harvested, the APCs, were given exogenous SIINFEKL peptide (1 nM) for 2 hours and kept at 37°C, in 5% CO_2_, and placed on a rotator to guarantee mixing. During this time, lymphocytes and splenocytes were harvested from an OT-1 mouse, and labeled with either CTV or CFSE according to the manufacture’s protocol (Molecular Probes). After the 2 hours of incubation, the B6 splenocytes were irradiated with 3,000 rad. Irradiated B6 splenocytes and CTV or CFSE stained OT-1 lymphocytes and splenocytes were mixed, 10 B6 derived cells per OT-1 derived cell, in sterile 96 well v-bottom plates. The inhibitors were then administrated and the plates were kept at 37°C and in 5% CO_2_ for 48 hours. After the 48 hours, cells were labeled with a fixable Live/Dead stain according to manufacture’s protocol (Molecular Probes), fixed in 2% PFA, and suspended in 90% MeOH at -20°C until staining for flow cytometry.

## Data analysis

Flow cytometry measurements were compensated and gated using FlowJo software. All other data analysis was performed using the scientific python software suite (SciPy), figures were produced in matplotlib [47], and Gaussian mixture modeling performed using scikit-learn [48].

## Acknowledgments

This research was funded by NIH R01 AI083408 and NIH U54 CA148967. A.E. is supported by the Human Frontier Science Program grant LT000123/2014. Furthermore, we thank Neal Rosen for commentary and for generously sharing inhibitors. We would also like to thank Jacqueline Bromberg for her valuable comments and discussions.

## Author Contributions

R.V. developed models; R.V. and G.A.-B. designed experiments; R.V. and A.E. analyzed data; R.V., A.E. and G.A.-B. performed experiments and wrote the manuscript.

## Additional Information

Supplementary Material includes model derivations, analysis, and supplementary data. Competing financial interests : The authors declare no competing financial interests.

